# Management of family relationship information for a three-generation cohort study

**DOI:** 10.1101/510073

**Authors:** Kazuro Shimokawa, Mami Ishikuro, Taku Obara, Hirohito Metoki, Satoshi Mizuno, Satoshi Nagaie, Masato Nagai, Chizuru Yamanaka, Hiroko Matsubara, Mayumi Kato, Yuki Sato, Soichi Ogishima, Takako Takai-Igarashi, Masahiro Kikuya, Atsushi Hozawa, Fuji Nagami, Shinichi Kuriyama, Kengo Kinoshita, Masayuki Yamamoto, Hiroshi Tanaka

## Abstract

A system for inputting and storing family information, named “BirThree Enrollment,” was developed to promote a birth and three-generation cohort study (BirThree Cohort Study), and this system was operated successfully. In the study, it was necessary to satisfy many operational demands. Input information is overwritten and changed continuously. Complex kinship information must be quickly and accurately input and corrected, and information on those families not yet recruited must be retrieved. For these purposes, many devices are needed, from an input interface to the internal data structure. In the field of genetic statistics, a simple standard expressive form is used for describing family structure. This form has sufficient information for genetics; however, we developed this form further for our purposes in conducting the BirThree Cohort Study. To provide information about family roles as required in the BirThree Cohort Study, we expanded the data structure, and constructed the system that is able to be used for the daily operation. In our system, family pedigree information is stored along with initial clinical information, and enabled the input of all self-reported information to the data base. Operators are able to input this family information before the day is out. As a result, when recruitment is completed, family information will be completed concurrently. Therefore, it is possible to immediately know a certain person’s family structure. By using our system, data correction was improved dramatically, and the system was operated successfully. This study is the first report of the method for storing three generations of family data.

## 1. INTRODUCTION

The Tohoku Medical Megabank (TMM) Project aims to provide creative reconstruction methods and solve medical problems arising from the Great East Japan Earthquake (GEJE), which occurred in March 2011 [1, 2]. In the TMM Project, two prospective cohort studies were initiated in Miyagi and Iwate Prefectures: a population-based adult cohort study called the TMM Community-Based Cohort Study (TMM CommCohort Study) and a birth and three-generation cohort study called the TMM Birth and Three-Generation Cohort Study (TMM BirThree Cohort Study) [3]. For the CommCohort Study, the TMM recruited participants at those sites where the specific health checkups of the annual community health examination were performed, at seven Community Support Centers in Miyagi Prefecture, and at five satellites in Iwate Prefecture. For the TMM BirThree Cohort Study, the TMM recruited pregnant women at obstetric clinics and hospitals in Miyagi and Iwate Prefectures along with their children and the children’s siblings, fathers (husbands), grandparents, and other family members. The TMM project contributed to the establishment of an integrated biobank that combines physiological and clinical data with genomics data. Additionally, some of the bio specimens collected by the TMM project were analyzed by other research laboratories, and stored in the TMM database, so as to facilitate a full range of omics analyses [4, 5].

In the CommCohort Study, many of the parties concerned were unrelated individuals. Thus, information on kinship relationships was not used for earlier ToMMo research, such as the 1KJP reference panel [6] that used the information collected for the CommCohort Study. They are the results of several preceding studies [7–9] that did not use family information. In contrast, all of the participants in the BirThree Cohort Study were related to other participants.

Birth cohort studies were among the first types of research to use data on family relationships. Preceding studies on birth cohorts include LifeLines, ALSPAC, MoBA, DNBC, and BiCCA [10–14]. Some of these studies successfully recruited 100,000 or more pregnant women in the birth cohorts. In these birth cohorts, a primary aim was to follow mothers and their newborns; therefore, recruitment of pregnant women is given priority. There were few opportunities to recruit additional relatives, and thus the recruitment and enrollment of other relatives was uncommon.

One of the key features of the BirThree Cohort Study is that three-generation cohorts were recruited, rather than just birth cohorts. Therefore, inputting and treating family information is more complex in a three-generation cohort study than in other cohort studies. The Lifelines study positively collects information not only on a pregnant woman and the child, but also on the father and other family members, as much as possible. Thus, it is an important initial research for treating family information. Such family information, however, is stored and maintained by a different system than that used for clinical information. As a result, massive data reduction after recruitment is indispensable, and much work might be needed to maintain the correspondence of clinical information and family information.

As a result, it was necessary to devise a data structure to operate the three-generation cohort recruitment. To express kinship information, a common data structure has traditionally been used in the field of statistical genetics [15–17]. Because the traditional data format can describe a parent-child kinship (Fig. 1A, B), it is necessary and sufficient for drawing family pedigrees and expressing genetic relationships. However, in some large-scale studies, such as the BirThree Cohort Study, this data format is insufficient to meet the various operational demands of three-generation cohort research. For instance, the format is not sufficient for the enrollment of relatives other than parent-child, such as grandfather-child. Moreover, there are some difficult problems concerning the data operation for the withdrawal of consent. This paper explains the basic idea underlying the BirThree Cohort Study and the data structure based on the specification. The specification is chiefly organized to handle the following operational matters: “Data description,”, “Retrieval,” “Consent withdrawal,”and “Family roles.”

**Fig. 1.**
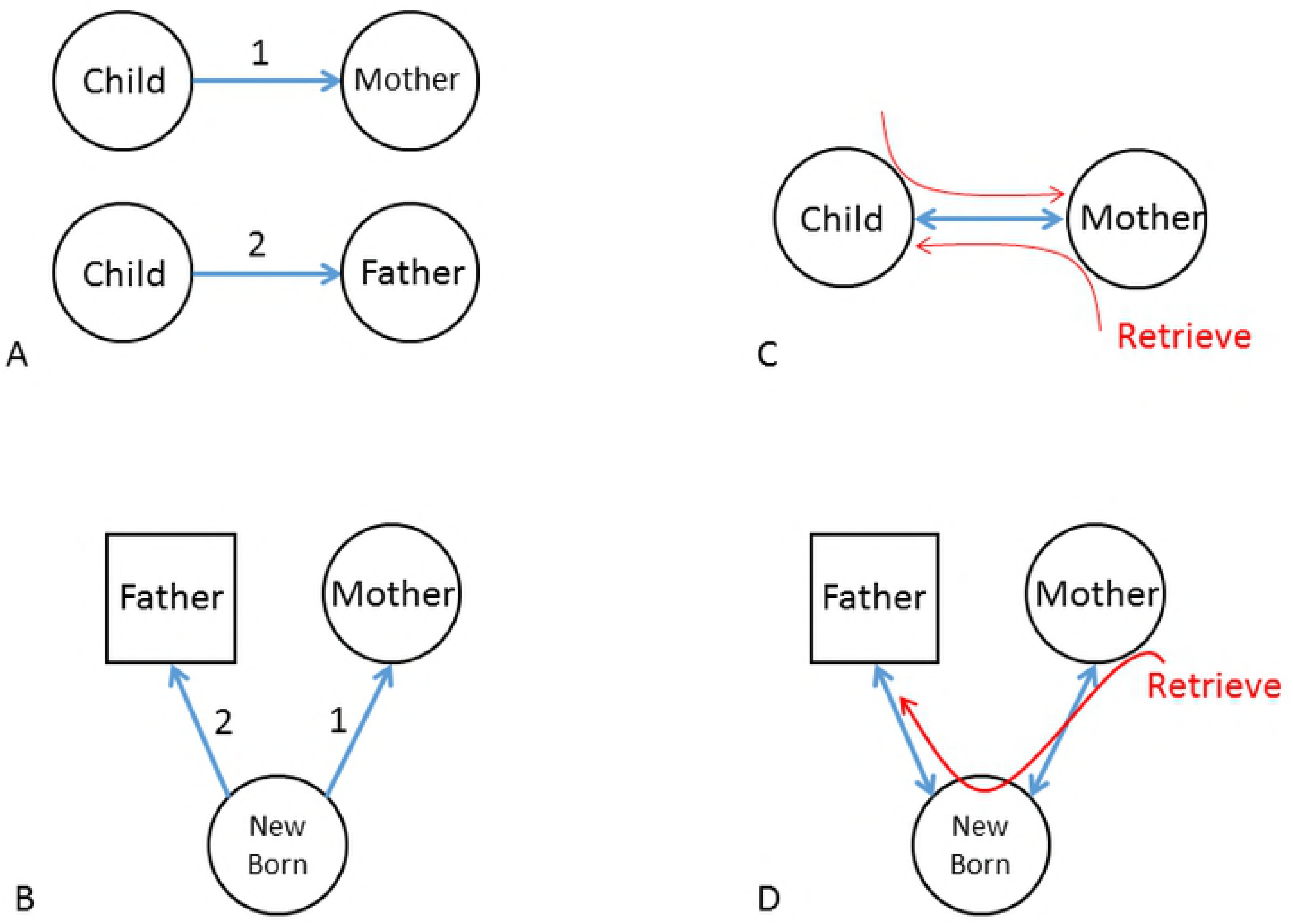
Basic Related Lines describing the relationship between two people. A and B. One-way Related Lines. The child cannot be traced from the mother (or father) by using this line. C. A Bidirectional Related Line, which can be used to trace from a child to a mother and from a m other to a child. D. We can identify the father by tracing the newborn (child) from the mother by using a Bidirectional Related Line.

## 2. METHODS AND RESULTS

### 2.1 Bidirectional Related Line

The concept of the data structure used in typical statistical genetics is described in Fig. 1A, B. Related Lines 1 and 2 show the relationships between the Child Identification Number (ID) and the Maternal ID or Paternal ID, respectively. These Lines are directional, such that the mother can be retrieved only by the certain child (Related Lines 1), and the child cannot be retrieved by their mother. This structure is used in the kinship2 [18] package in the R statistics program and suffices to describe genetic relationships. There are some problems, however, with retrieving family relationships by using the Related Lines. For instance, to find the child of a certain mother, it is necessary to search all of the children in the database in the worst case scenario. Thus, we first made the Related Line bidirectional (Fig. 1C). Each related line can thus be defined by two kinds of edges (ex., Child to Mother, Mother to Child) in a direct acyclic graph, rather than one edge in an undirected graph, following a basic idea of network theory [19–21]. With this new bidirectional line, it becomes easier to trace father from mother, by retrieving from mother to child, and from child to father. In this way, relationships between parents are retrieved more quickly (Figs. 1B, D). Retrieving all members in the family becomes possible by using this line. However, retrieval might become difficult when there is a member who is not participating in the family. We discuss this problem in the following paragraph.

### 2.2 Enrollment System of three generations

#### 2.2.1 Extended Line

The necessity for registering the person who doesn’t have a parent-child relationship appears, while registering a family’s member. One example is that of the relationship between two participants who are connected by a nonparticipant, which is not expressible (Fig. 2A). In this case, the relationship between “Newborn” and “Grandfather” cannot be made without the father’s participation. One solution to this problem is to define another type of Related Line (the Extended Related Line). The Extended Related line connects the relationship from any person in a seven-member family to the other person. Therefore, this line can connect two members in the seven-member family who do not have a parent-child relationship. This approach is thought to work well when the scale of the cohort is small and the lineage is simple. We explain the concept of defining an Extended Related Line for registration below.

**Fig. 2.**
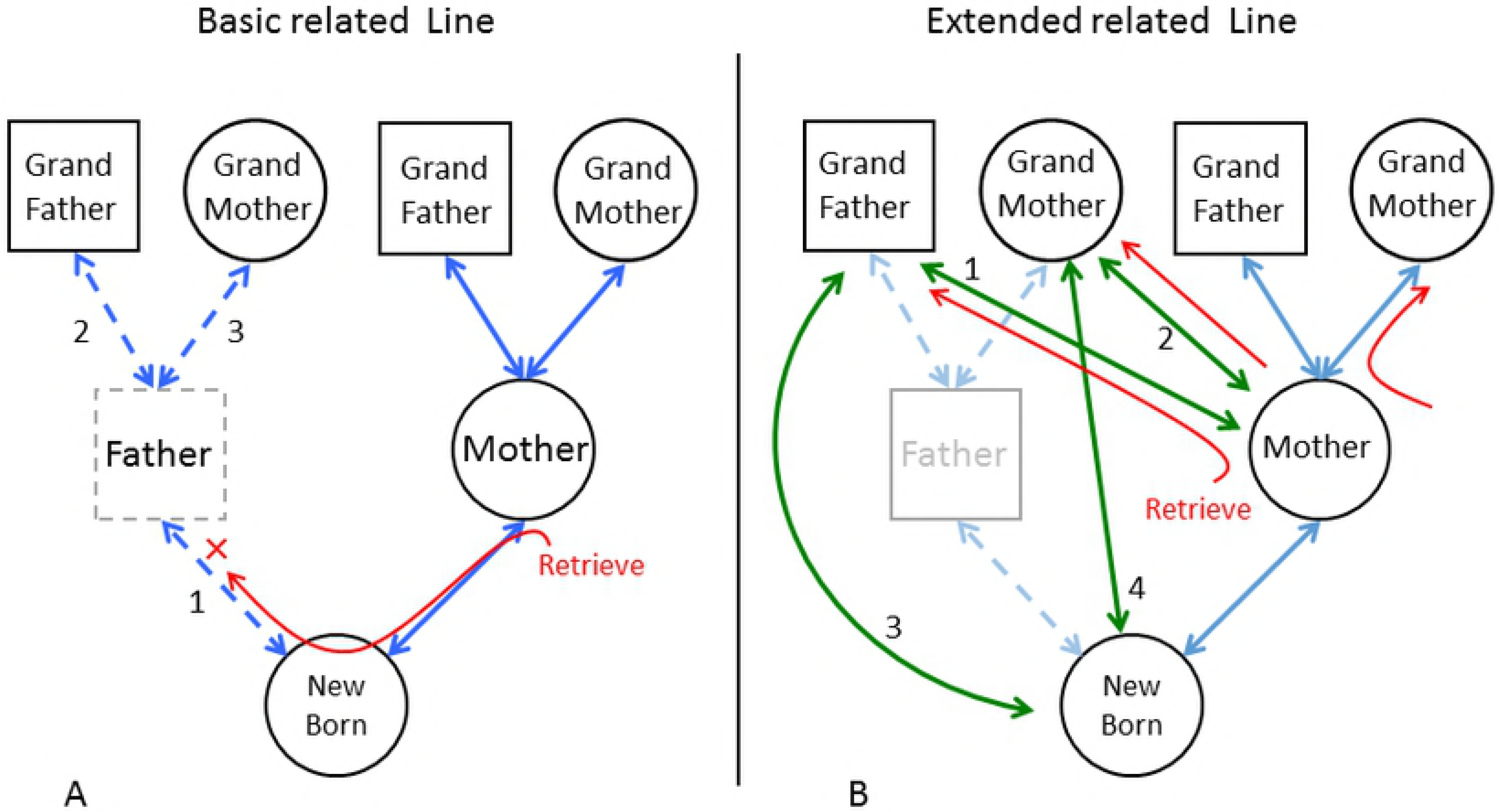
Basic and Extended Related Lines. A. A Basic Related Line only connects the child with the parents. The dotted rectangle shows the family member(s), who have not yet been recruited. When the father is a nonparticipant, a related line is not connected with grandparents by the newborn baby. B. An Extended Related Line expresses the relationships among seven family members using many predefined lines. Four extended related lines (green arrow) are selected from among the 21 extended significantly related lines to enroll this family.

According to the inclusion criteria for the TMM BirThree Cohort, pregnant women must be recruited first in a three-generation family. Then, other family members in the three-generation cohort (newborn (child), father, grandmother, grandfather, grandmother-in-law, and grandfather-in-law) are recruited. These seven family members (the pregnant woman and the other 6 family members) are recruited as one unit. Thus, when attention is paid to a specific participant, the participation status of the other six members should be understandable. By means of the bidirectional Related Line, a mother related to a child and the child’s father can be retrieved if all three people are participating, as shown in Fig. 1D. It is impossible, however, to find another family member who is connected through a nonparticipant. For instance, when the husband is not a participant, the grandfather-in-law (paternal grandparent) cannot be enrolled and retrieved from the mother’s information by using the bidirectional Related Line (see the red arrow in Fig. 2A).

One solution would be to extend the Related Line to obtain an Extended Related Line. The green arrows in Fig. 2B show this type of Extended Related Line. Operators can easily find members of the family connected to the pregnant woman and Newborn by using an Extended Related Line. The Extended Related Line offers a method for defining all of the relationships in the cohort. Therefore, the example problem can be solved by defining Extended Related Lines between the mother and grandfather-in-law (green arrow 1), the mother and the grandmother-in-law (green arrow 2), the Newborn and the grandfather-in-law (green arrow 3), and the Newborn and the grandmother-in-law (green arrow 4) (see Fig. 2B). The relationships between these individuals can be enrolled and retrieved from the mother (a pregnant woman) to the grandparents, or from the Newborn to the grandparents, through Extended Related Lines (Fig. 2B), even if the father does not participate in the cohort. Table 1 presents the number of Extended Related Lines needed for all patterns of seven family members.

**Table 1.**
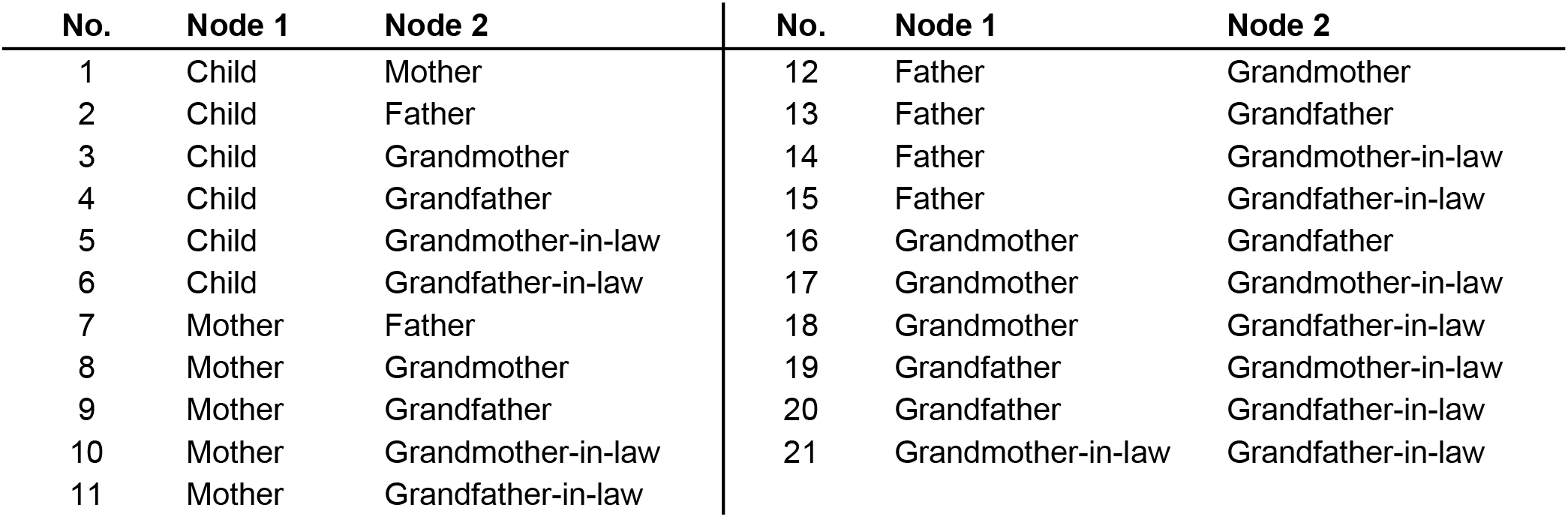
Example of an Extended Related Line

| No. | Node 1 | Node 2 | No. | Node 1 | Node 2 |
| --- | --- | --- | --- | --- | --- |
| 1 | Child | Mother | 12 | Father | Grandmother |
| 2 | Child | Father | 13 | Father | Grandfather |
| 3 | Child | Grandmother | 14 | Father | Grandmother-in-law |
| 4 | Child | Grandfather | 15 | Father | Grandfather-in-law |
| 5 | Child | Grandmother-in-law | 16 | Grandmother | Grandfather |
| 6 | Child | Grandfather-in-law | 17 | Grandmother | Grandmother-in-law |
| 7 | Mother | Father | 18 | Grandmother | Grandfather-in-law |
| 8 | Mother | Grandmother | 19 | Grandfather | Grandmother-in-law |
| 9 | Mother | Grandfather | 20 | Grandfather | Grandfather-in-law |
| 10 | Mother | Grandmother-in-law | 21 | Grandmother-in-law | Grandfather-in-law |
| 11 | Mother | Grandfather-in-law |  |  |  |

This method has operational problems, however, especially during registration. It is necessary to define many Related Lines, and operators should correctly select the proper Related Lines when enrolling each participant. Even in a case such as the one shown in Fig. 2B, which is not very complex, four (2 x 2 (bidirectional)) new different lines are needed to make connections from the mother. This solution might work when the data input operator is able to spend sufficient time or when complex family relationships need not be drawn. In the case, however, in order to connect seven family members with Related Lines, it is necessary to select the correct colored lines from among 21 x 2 (bidirectional) = 42 types of Related Lines (see Table 1). This operation becomes a considerably time-consuming load for the operator. During the first stage of recruitment of the BirThree Cohort, this idea was adopted for our system, and put into operation. However, operation of the system becomes difficult, as the scale of the BirThree Cohort in the TMM became large-scale. Therefore, this idea was not adopted in the present system, although we introduce the idea here for reference.

When a Related Line connects every set of seven members, 21 x 2 (bidirectional) different Related Lines are needed.

#### 2.2.2 BirThree Enrollment system

We have developed the BirThree Enrollment system instead of using the system of Extended Related Lines in order to accurately describe complex family roles and avoid the input of incorrect data. It is thought that such a family input system is indispensable to recruit the family members, and this system will become the main current in the Cohort study in the future. The BirThree Enrollment system has a family role table that is used to enroll seven family members. This idea considers two critical factors. One is to apply a pregnant ID, instead of a family ID, to manage the family role table. A pregnant ID is allocated at each pregnancy. A pregnant woman and the number of pregnancies are always identified by a pregnant ID, and the ID is a key to each family role table. The family role table stores seven family member’s IDs and their roles. It corresponds to the red or yellow rectangle in Figure 3. By using the pregnant ID, it is possible to recruit members over a long term with stability. On the other hand, a family ID is an idea that comes from the field of genomic analysis. The participants who exist in the same family tree (two or more pregnant women can be included) will have same family ID. It is difficult to use this idea with the Cohort enrolling system. This is because the family can be mutually connected after a long period of recruitment, and the number of families IDs decreases. Then, the pregnant woman as a proband in the family cannot be identified.

**Fig. 3.**
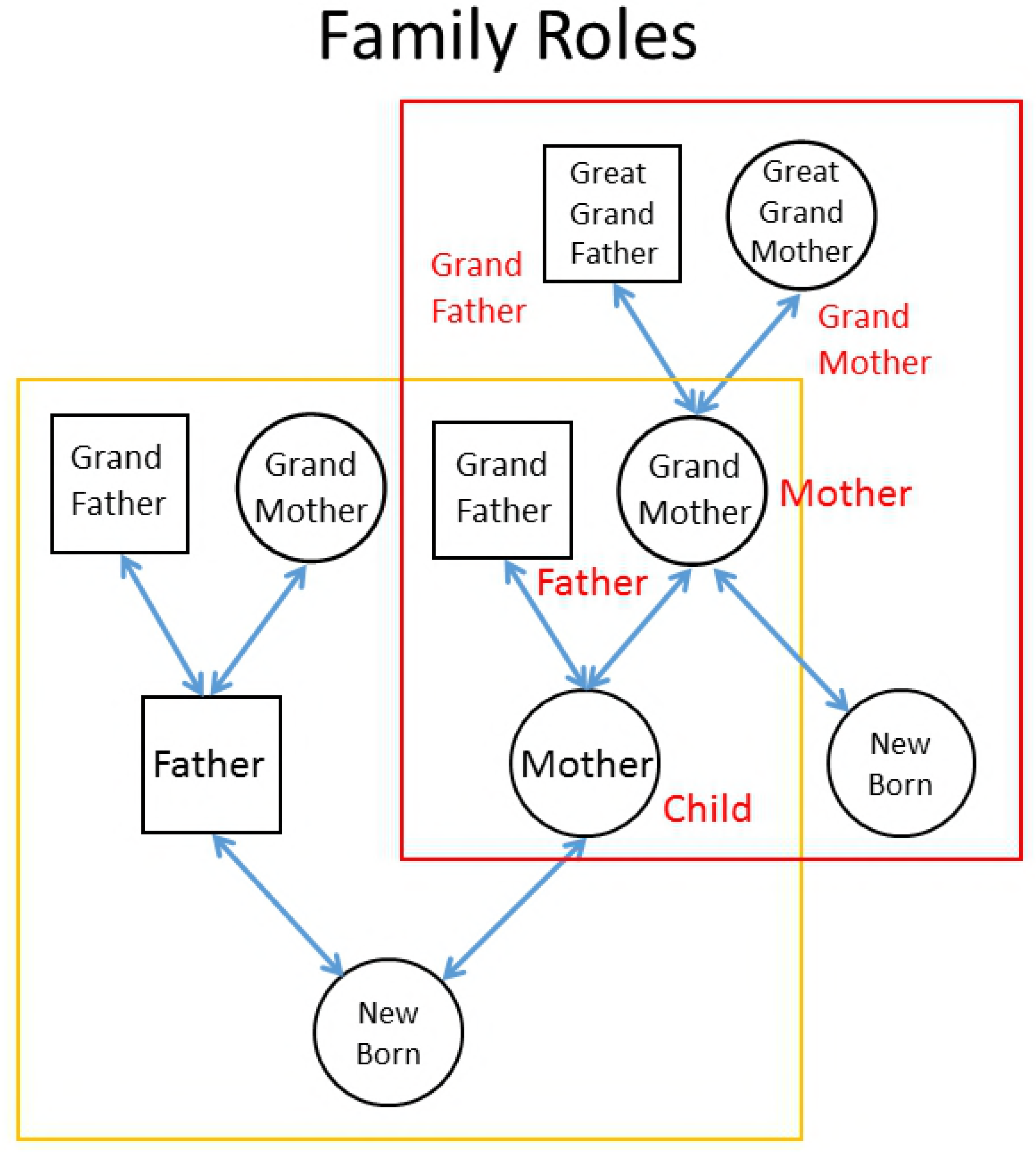
Explanation of family role tables when a participant has two or more family roles. In this figure, the “mother” in the family role table enclosed within a red rectangle is also a “grandmother” in the family role table within a purple rectangle. When two or more family role tables are applied for one person, it means that the individual has different family roles in each table.

The other factor is the family information batch entry screen, a new family data enrolling method (Fig. 4). The batch entry screen makes it possible to design a comprehensible data registration interface (see supplemental file). The entry screen not only enables the registration of relationships through nonparticipants, but also facilitates later retrieval and eases registration. A family role table is newly prepared when the pregnant woman is enrolled. At that time, the other six family members are added as provisional participants for whom recruitment is necessary in an empty column (see Section 2.4 Fig. 4). When some of these six non-participants are recruited, and some empty columns are filled, related lines are automatically drawn by the BirThree Enrollment system.

**Fig. 4.**
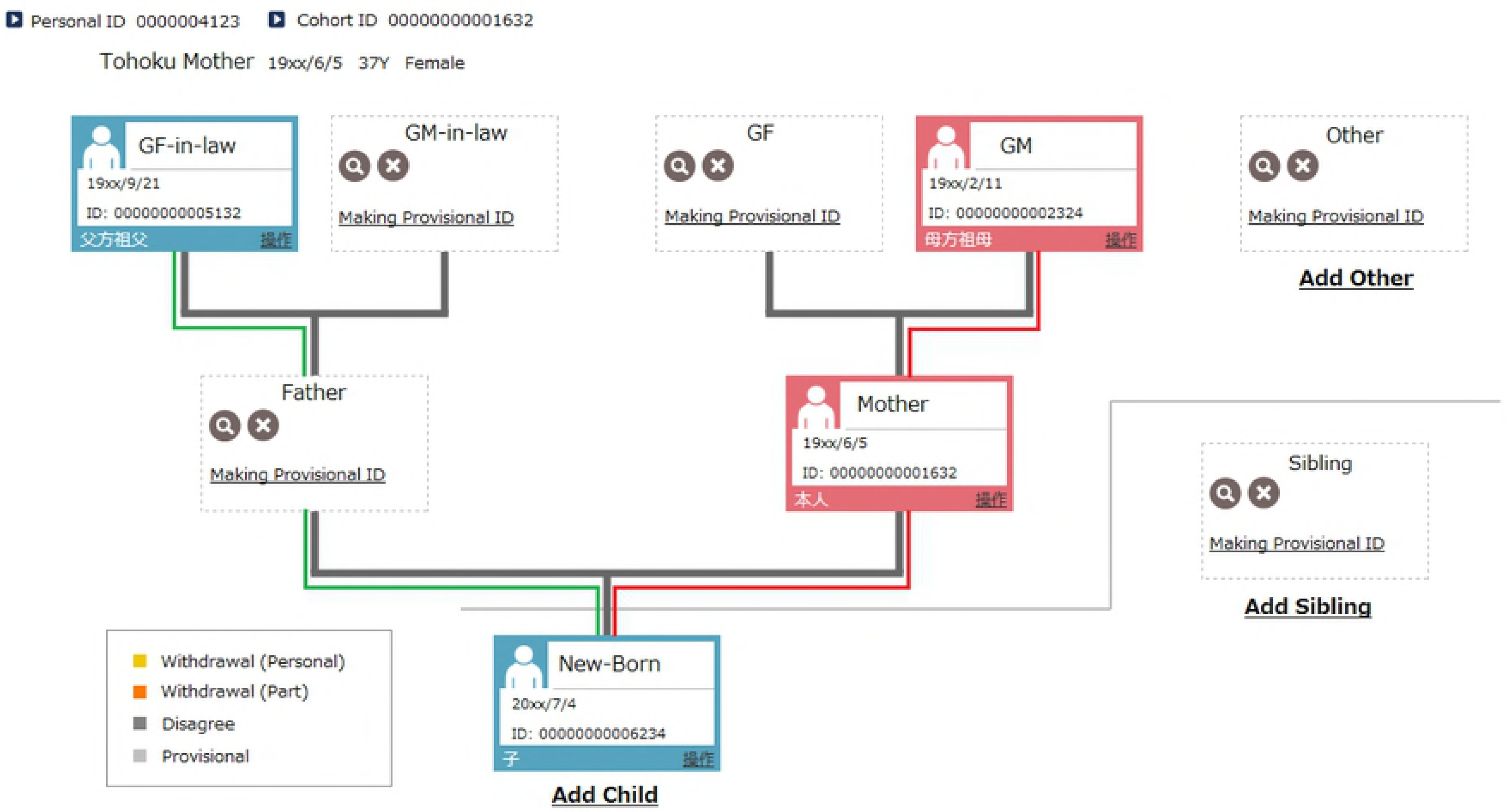
Actual family information batch entry screen. The system is designed so that each family member is enrolled in a specific position in the screen. At the same time, when a member is enrolled, he or she is also enrolled in the family role table, and the Related Lines between members are automatically connected. The dotted rectangle shows other members of the family who have not yet been recruited. In this figure, “Provisional ID” refers to a temporary ID and indicates that the member has not yet been recruited. Related Lines that correspond to the red line in this figure have already been connected, because the three members (GM (Grand Mother), Mother, New-Born) were already recruited. When “father” is enrolled, the Related Lines that correspond to the green line will be automatically connected. When “father” withdraws after participation, the Related Lines corresponding to the green line are kept (not deleted). The number of family members can be increased by clicking the “Add sibling” or “Add Other” button (See the supplementary file).

As in the example with the nonparticipating father shown in Fig. 2A, even in the situation whereby a pregnant woman and her father-in-law cannot be connected directly, the Related Lines 1, 2, and 3 in Fig. 2A among the family members are automatically drawn by our BirThree Enrollment system, and this father is automatically registered as a provisional member. Therefore, lines are always connected whenever family members are retrieved.

The family information batch entry screen allows one person to be enrolled in two or more role tables at the same time. As a result, one person can have two different roles in two different families. Fig. 3 shows a person enrolled as a mother in the family role table enclosed by the red rectangle and as a grandmother in the family role table with the purple rectangle. Such complex expressions are possible in this system. The BirThree Enrollment system was adopted because of this ability.

### 2.3 Implementation

Fig. 4 shows the family information batch entry screen that was used for recruitment in the TMM BirThree Cohort Study. In the implementation, the pregnant woman’s ID is treated as the main key in the family role table. Therefore, the pregnant woman is always in the family role table. This entry screen is an enrolling screen that is used by the operator during recruitment. All of the positions that the seven family members occupy are created in advance.

The family role table corresponds to the entry screen. The input system registers data in both the family role table and the Related Lines. Related Lines are automatically formed for unit members (pregnant woman (mother), newborn (child), father, grandmother, grandfather, grandmother-in-law, and grandfather-in-law). Uncles, aunts, and cousins are also other members of the unit; therefore, operators must draw those Related Lines by hand.

While retrieving the family structure, the database extracts family information by reading only the table of the corresponding family role. Tracing a related line for the retrieval is not necessary. Therefore, the load on the system is minimized. Moreover, it becomes easy to call a participant through the pregnant woman, because the system displays whether there is a family member who has not yet been recruited during retrieval.

### 2.4 Withdrawal of Consent

When a pregnant woman withdraws her consent, the consent of her new born child is withdrawn. On the other hand, other family members (siblings, father, Grandparents) stay as participants in principle, because their consent forms are still effective. Alternatively, they are registered as a participant in the follow-up survey by the TMM Project. In contrast, when a member other than a pregnant woman withdraws consent, only that person’s name and information are deleted. For that case, the Related Line containing that person and other members of the family is not deleted, and the family role table remains. For example, when a participating father withdraws his consent, the Related Lines 1, 2, and 3 in Fig. 2A are maintained to avoid re-recruiting people who have been withdrawn.

An important point of our system is to process the consent withdrawal in real time. A problem might not occur to process the consent withdrawal after recruitment ends, when the recruitment period is short. When the system continues working for a long time, however, the consent withdrawal process needs to be reflected at once. This is because when the follow-up survey and re-recruiting is performed, withdrawal information must reflect the process up until then.

Such a mechanism must be carefully implemented to ensure that it does not contradict the contents written in the informed consent and withdrawal documents. This is because the level of information that should be removed is written into the withdrawal document. It is necessary to design the informed consent and withdrawal documents so that no contradictions should occur in data processing.

### 2.5 Error Correction

We designed a function to identify and report discrepancies in the database. This function was designed for important uses and had a significant influence on the quality of the cohort information. We had to report discrepancies immediately when mistakes in the registration data caused severe problems. Here, we explain the check function, which was used for information about family relationships.

Bidirectional Related Lines are important items to check, because they include information that overlaps among family role tables. The BirThree Enrollment system automatically registers all of the Related Lines for the seven family members. Therefore, family role tables and those related lines never contradict each other when all the family information is entered through the family information batch entry screen. Not all relationships, however, can be entered by using the family information batch entry screen. For instance, the Related Lines between a pregnant mother and her siblings, and between a newborn and its siblings are drawn by hand. The data correction routine recomposes the family relationships by using the Related Line, and compares it with the information in the family role table. Any discrepancies are reported to the data entry operators for correction. Due to this routine work, errors in family relationships were minimized. Related Lines and family role tables overlap; therefore, they were thoroughly tested for contradictions. This report has been confirmed by the data entry operators, and the data correction routine operates correctly every day.

## 3 DISCUSSION AND CONCLUSIONS

We created an input and data registration system, “BirThree Enrollment,” for a birth and three-generation cohort study and successfully collected data from more than 70,000 BirThree Cohort participants, which were required for a research platform [10, 22–25]. This system was used by more than 150 BirThree Cohort Genome Medical Research Coordinators (GMRC), and development was advanced to the fifth version.

TMM’s work is the first attempt in the world to create a three-generation Cohort Study that collects participant data on a 100,000-person scale. Our study was able to achieve the large-scale number of participants, although some other birth cohort studies were unable to collect the aimed numbers of participants, and closed [26]. The operation of this system contributed to the success of the research. From the viewpoint of data science, this work is a method for displaying and browsing a 7-member family tree, and provides the data structure required to achieve it. It is practically effective even when a family’s information changes dynamically by withdrawal of the agreement.

The remaining problems include handling information on a participant who gave birth two times or more and a father who divorced the pregnant woman. Our system was insufficient to decide which data to store for the person who had participated two times or more. The other problem, the “father who divorced,” did not actually occur for pregnant women who participated two or more times with a different husband for each pregnancy. It was unnecessary to connect information about the different fathers.

It is difficult to complete a three-generation cohort project with high efficiency, because the process of inputting family information is complicated. To address this problem, computational support is extremely important. In this study, many problems involved with treating family information have been solved with reference to six viewpoints. We expect this study to be useful for the next third-generation cohort study.

## 4. ACKNOWLEDGMENTS

This study was supported by grants from the Reconstruction Agency, the Ministry of Education, Culture, Sports, Science and Technology (MEXT), and the Japan Agency for Medical Research and Development (AMED). This study was also supported by JSPS KAKENHI (grant number JP16K12717).

The authors express their gratitude to the people of Japan and of the world for their valuable support of the areas affected by the GEJE after the disaster. We thank the members of ToMMo and IMM, including GMRCs and the office and administrative personnel for their assistance. We also thank Mr. Hiroshi Hanzawa, Mrs. Tomomi H. Oonuma, Mr. Junji Takada, Mrs. Yuka O. Takahashi, and Mr. Gosuke Takahashi for technical assistance.

